# Expanding Interdisciplinarity: A bibliometric study of medical education using the MEJ-24

**DOI:** 10.1101/2023.03.22.533841

**Authors:** Lauren A. Maggio, Joseph A. Costello, Anton B. Ninkov, Jason R. Frank, Anthony R. Artino

**Author notes:** Correspondence should be addressed to Lauren A. Maggio, Department of Medicine, Uniformed Services University of the Health Sciences, 4301 Jones Bridge Rd., Bethesda, MD 20814. @LaurenMaggio. @LaurenMaggio. @jojo_costello. @BoudreauNinkov. @drjfrank. @mededdoc.

## Abstract

**Introduction:** Interdisciplinary research has been deemed to be critical in solving society’s wicked problems, including those relevant to medical education. Medical education research has been assumed to be interdisciplinary. However, researchers have questioned this assumption. The present study, a conceptual replication, provides an analysis using a larger dataset and bibliometric methods to bring more clarity to our understanding on the nature of medical education interdisciplinarity or lack thereof.

**Method:** The authors retrieved the cited references of all published articles in 24 medical education journals between 2001-2020 from the Web of Science (WoS). We then identified the WoS classifications for the journals of each cited reference.

**Results:** The 24 journals published 31,283 articles referencing 723,683 publications. We identified 493,973 (68.3%) of those cited references in 6,618 journals representing 242 categories, which represents 94% of all WoS categories. Close to half of all citations were categorized as “education, scientific disciplines” and “healthcare sciences and services”. Over the two decades studied, we observed consistent growth in the number of references in other categories, such as education, educational research, and nursing. Additionally, the variety of categories represented has also increased from 182 to 233 to include a diversity of topics such as business, management, and linguistics.

**Discussion:** This study corroborates prior work while also expanding it. Medical education research is built upon a limited range of fields referenced. Yet, the growth in categories over time and the ongoing increased diversity of included categories suggests interdisciplinarity that until now has yet to be recognized and represents a changing story.

## Introduction

Medical education research has been described as interdisciplinary with its researchers hailing from a variety of fields and disciplines (e.g., clinical medicine, education, sociology) and drawing upon a variety of methods, methodologies, and epistemic traditions [1,2]. Policy makers, funders, and scientists have deemed interdisciplinary research as crucial [3] and requisite for helping society solve its wicked problems; i.e., problems that are complex in scope and lack clear solutions [4]. Medical Education has its share of wicked problems across a range of topics from curricular reform to health systems leadership [5-7]. This broad range of wicked problems suggests that interdisciplinarity is critical for solving medical education’s challenges. However, in two recent studies, Albert and colleagues questioned the interdisciplinary nature of medical education research, concluding that the characterization of medical education as an interdisciplinary field is unsupported by evidence [8,9]. While these studies are informative, we do not believe they tell the entire story due to their limited size and scope. To address the limitations of this prior work, we conducted a conceptual replication of one of Albert et al.’s studies [8] with a much-expanded data set to provide a more complete description of the interdisciplinary nature of medical education research.

The National Academy of Science defines interdisciplinary research as: “research by teams or individuals that integrates information, data, techniques, tools, perspectives, concepts, and/or theories from two or more disciplines or bodies of specialized knowledge to advance fundamental understanding or to solve problems whose solutions are beyond the scope of a single discipline or field of research practice” [10, p. 27]. For over two decades, researchers have studied interdisciplinarity using bibliometrics, which is the examination of publications and publication metrics to understand publishing trends, including relationships between papers through citation patterns [11]. For example, Lariviere and colleagues studied the interdisciplinarity of articles indexed in the database Web of Science over a 110-year period [12]. In this work, the authors defined interdisciplinarity of a particular article as the percentage of its cited references published in journals that are outside its discipline, which they referred to as “citations outside of category” [12]. The authors found that over time authors have increasingly cited articles from outside their discipline thus becoming more interdisciplinary across the century.

In 2020, Albert and colleagues drew on Lariviere’s work to study the “widespread assumption” that medical education research is interdisciplinary [8]. The authors studied 64 research articles published in 2017 that were drawn from five medical education journals. Journals were selected because they had the highest impact factors for medical education journals at the time of the study. The selected articles cited 1,412 references, which the authors classified into six discipline-focused clusters. Based on the classification of the cited references, the authors found that 81% of the cited references were published in either medical education or clinical and health services journals. The authors concluded that medical education research stands predominantly on the foundation of these two domains. Cited references outside of these two closely related clusters, which would be considered citations outside of category, were in the minority (i.e., only 19%). In a follow-on study using the same data set, the authors compared the reference patterns of the medical education articles with a set of articles drawn from higher education [9]. Based on this comparison, the authors concluded that medical education research is “inwardly focused” in general and when compared to higher education. While Albert and colleague’s work is commendable, their sample is narrowly defined and represents a very small portion of published medical education articles. For example, between 2000-2020, 37,263 articles were published in 24 medical education journals [13]. This suggests that the 64 research articles, which represent only 0.17% of this entire sample of published articles, likely fail to fairly represent the field of medical education. Additionally, the focus on five journals, based on impact factor alone, does not take into account the range of medical education journals, including those that are newer to the field and may have a low or no impact factor. We believe such journals should not be discounted, especially if we consider the many well-documented limitations of impact factor as a measure of quality [14, 15].

To expand on Albert’s work, we conceptually replicate his work in this study by characterizing a much larger sample of cited references over two decades. In doing so, we hope to bring additional evidence to bear on the question of whether or not medical education research is interdisciplinary. We also hope to better identify potential trends that have occurred over the time period examined.

## Method

We analyzed the references cited by medical education articles published in 24 journals between 2001-2020. The large-scale nature of our data set necessitated that we diverge from the original study. As such, we did not conduct a direct replication, but instead performed a conceptual replication, which includes purposefully diverging from the earlier study’s methods [16, 17]. When conducting a conceptual replication, researchers seek to re-test or confirm a theoretical idea or hypothesis by examining different populations or using different study methods/measures in order to increase confidence in previous findings [18].

In this study, we diverged from Albert by including data from articles of all publication types (e.g., research articles, perspectives, reviews) published over a 20-year period in 19 additional medical education journals. Additionally, we leveraged Web of Science’s (WoS) standardized categories, which are applied by the platform’s trained indexers and based on a journal’s scope note and editor’s suggestion. This change enabled us to identify characteristics of cited references at scale using an automated approach. In Albert’s study, the authors used six self-created categories that were applied by the individual researchers, which we felt was unfeasible at scale and did not take advantage of the available standardized categories that have been selected by the editors of the journals.

### Data Collection

We included articles and their references published in the 24 journals featured in the MEJ-24 [19]. The MEJ-24 is a list of medical education journals that was created using bibliometric co-citation and has been proposed as a “seed set” of journals to be used by researchers conducting bibliometric research as a proxy for the field of medical education (See Zenodo [20] for a listing of all MEJ-24 journals and their date coverage in the data set. See Maggio [19] for details of the MEJ-24 creation). On August 27, 2021, we obtained WoS metadata for these source articles, including for each article the references that they cited. WoS is a bibliometric database that is composed of indexes. In this study we focused specifically on the WoS Core Collection, which includes four major indexes: the Science Citation Index Expanded (SCIE), Social Science Citation Index (SSCI), Arts & Humanities Citation Index (AHCI), and Emerging Sources Citation Index (ESCI). Two MEJ-24 journals, the *Journal of Graduate Medical Education* and the *Canadian Journal of Medical Education* are not indexed in the WoS Core Collection, thus we were unable to obtain relevant metadata and therefore these two journals are excluded from further analysis. We used WoS as it provides well-defined metadata and is the standard bibliometric tool used by bibliometricians [21]. We organized the data in an Excel spreadsheet.

To characterize the cited references, we relied on WoS Categories. WoS Categories represent a standardized subject categorization scheme for all journals in the WoS Core Collection. For example, the journal *Perspectives on Medical Education* is described by the two WoS Categories: *Education, Scientific Disciplines* and *Health Care Sciences and Services*. It is worth noting that there are 256 WoS categories ranging from acoustics to zoology. As demonstrated in the previous example, journals can and often are characterized by more than one WoS Category. In some cases, a journal has multiple WoS Categories because it is contained in more than one WoS index. For example, *Advances in Health Sciences Education* is in both SCIE and SSCI. In the SCIE it is described by the WoS Categories: *Health Care Sciences & Services* and *Education Scientific Disciplines* and then, in SSCI it is described as *Education & Educational Research*.

Thus, *Advances in Health Sciences Education* is described by three total WoS categories. (See Table 1 for additional examples).

**Table 1:**
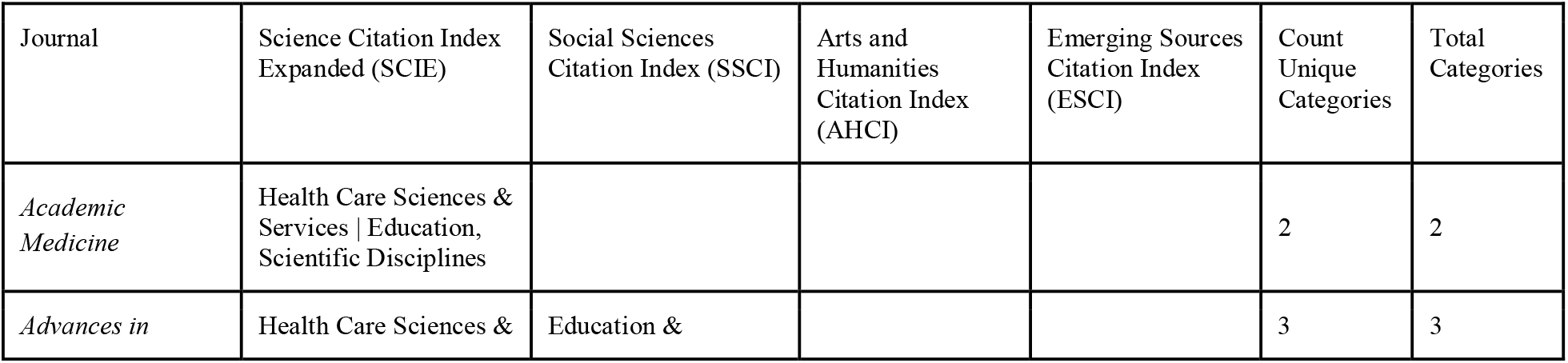
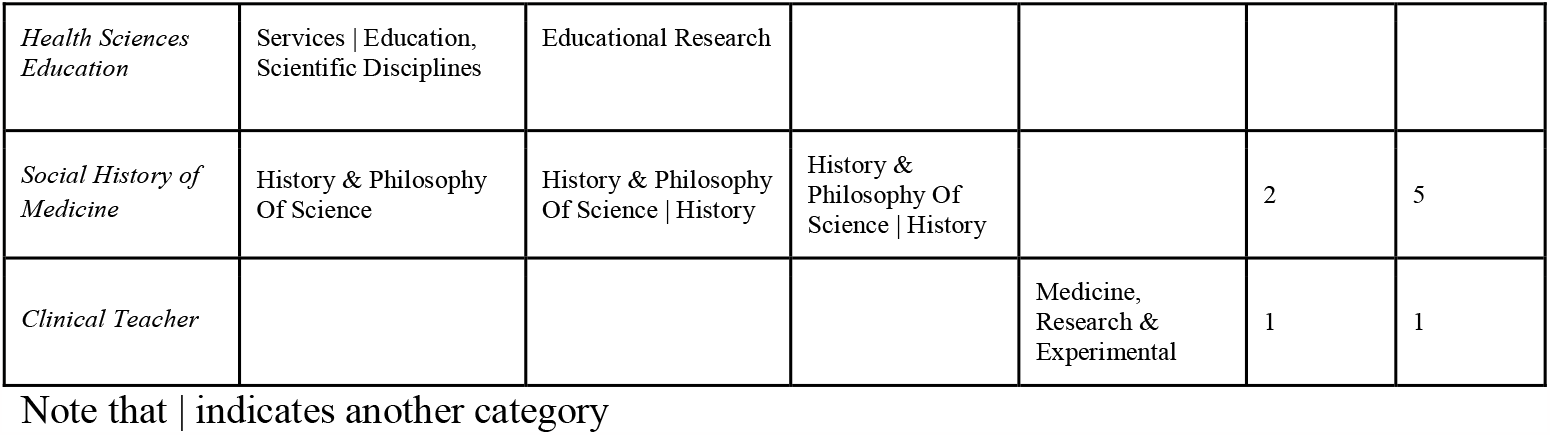
Examples of how journals are described by categories across Web of Science indexes

To sum the cited references and their categories, we wrote an Excel formula to count the appearance of a journal (see Zenodo [20] for additional details). As cited references are contained in journals that may be described using more than one WoS category, we decided to count each category equally. For example, there were 65,473 references to *Academic Medicine. Academic Medicine* is included in the WoS categories *Health Care Sciences and Services* and *Education, Scientific Disciplines*. Thus, in the instance of *Academic Medicine* we counted 65,473 references for each of the two WoS categories. Lastly, to characterize journals as clinical, we used the specialty classifications from the American Board of Medical Specialties [22] and consulted with JF, a clinician.

## Results

Of the 36,293 source articles in our data set, 5,010 (13.8%) did not have reference data because either the authors did not cite references (e.g., a letter to the editor, acknowledgement of reviewers) or the articles were published in CMEJ (n=405) or JGME (n=1947), the two journals not available in WoS. Thus, our analysis focuses on the 31,283 source articles for which we retrieved reference data. These articles cited 723,683 references. On average, articles had 23.1 references (SD= 20.6; MED=19). We identified 493,973 (68.3%) of those cited references in 6,618 journals with 242 categories, which represents 94% of all WoS categories (n=258) and 32% of all journals included in WoS. The 31.7% of missing cited references are likely alternative publication sources, such as websites, books, reports, etc., or they are references to journals that are not indexed in WoS. We focus our analysis on the 493,973 cited references.

Based on the count of cited references, *Academic Medicine* (n=65,473 references, 13.3%), *Medical Education* (n=50,372, 10.2%), and *Medical Teacher* (n=33,823, 6.8%) were the most cited journals. Outside of the MEJ-24, *JAMA* was the fourth most cited (n=14,473, 2.9%) followed by *Journal of General Internal Medicine* (n=9,317, 1.9%) and *Anatomical Sciences Education* (n=7,612, 1.5%) ranked as seventh and eighth, respectively. See Table 2 for the top 10 journals based on count of cited reference appearances. The journals in the MEJ-24 all appeared in the top 1000 journals with *Academic Medicine* (n=65,473) being the most cited and *Focus on Health Professions Education* (n=39) the least. See Zenodo [20] for a listing of MEJ-24 journals by count of cited reference appearances. Of the 22 journals in the MEJ-24 - with metadata we could analyze - they accounted for 201,521 (40.8%) of cited references.

**Table 2:** Top 10 journals based on Count of Cited References (n=493,973)

| Journal | Count of source articles (%) | Count of cited references (%) |
| --- | --- | --- |
| *Academic Medicine* | 6428; 20.5 | 65473; 13.3 |
| *Medical Education* | 4369; 14.0 | 50372; 10.2 |
| *Medical Teacher* | 4735; 15.1 | 33823; 6.8 |
| *JAMA* |  | 14473; 2.9 |
| *Advances in Health Sciences Education* | 1154; 3.7 | 9839; 2.0 |
| *BMC Medical Education* | 3033; 9.7 | 9572; 1.9 |
| *Journal of General Internal Medicine* |  | 9317; 1.9 |
| *Anatomical Sciences Education* | 868; 2.8 | 7612; 1.5 |
| *The New England Journal of Medicine* |  | 7094; 1.4 |
| *Teaching and Learning in Medicine* | 1071; 3.4 | 6386; 1.3 |

Cited references hailed from journals present across all WoS Indexes with SCIE (n=3519 journals), SSCI (n=2229), and ESCI (n=1274) most prevalent. Only 319 journals were present in the AHCI. Seven hundred and three journals were contained in more than one index with the most common pairing of indexes being SCIE and SSCI (n=544). (See Zenodo [20] for a count of journals in all indexes).

Cited references were published in 6,618 journals and described by 242 WoS categories of which we considered 33 to be clinically focused (e.g., Anesthesiology, Microbiology, Pediatrics). Journals were assigned by WoS between 1-6 categories, with an average of 1.5 categories.

We provide counts of categories at the journal level and cited reference level. At the journal level, each journal was counted only once per category. For example, although there are many cited references published in *Medical Education*, for this journal its categories of Education, Scientific Disciplines and Healthcare Sciences and Services are counted once. At the journal level, the most common categories were *Education and Educational Research* (n=460 journals), *Public, Environmental and Occupational Health* (n=383 journals), and *Nursing* (n=283 journals). See Figure 1 for the top 25 categories. We identified 2,239 journals that we categorized as clinically-focused journals.

**Figure 1:**
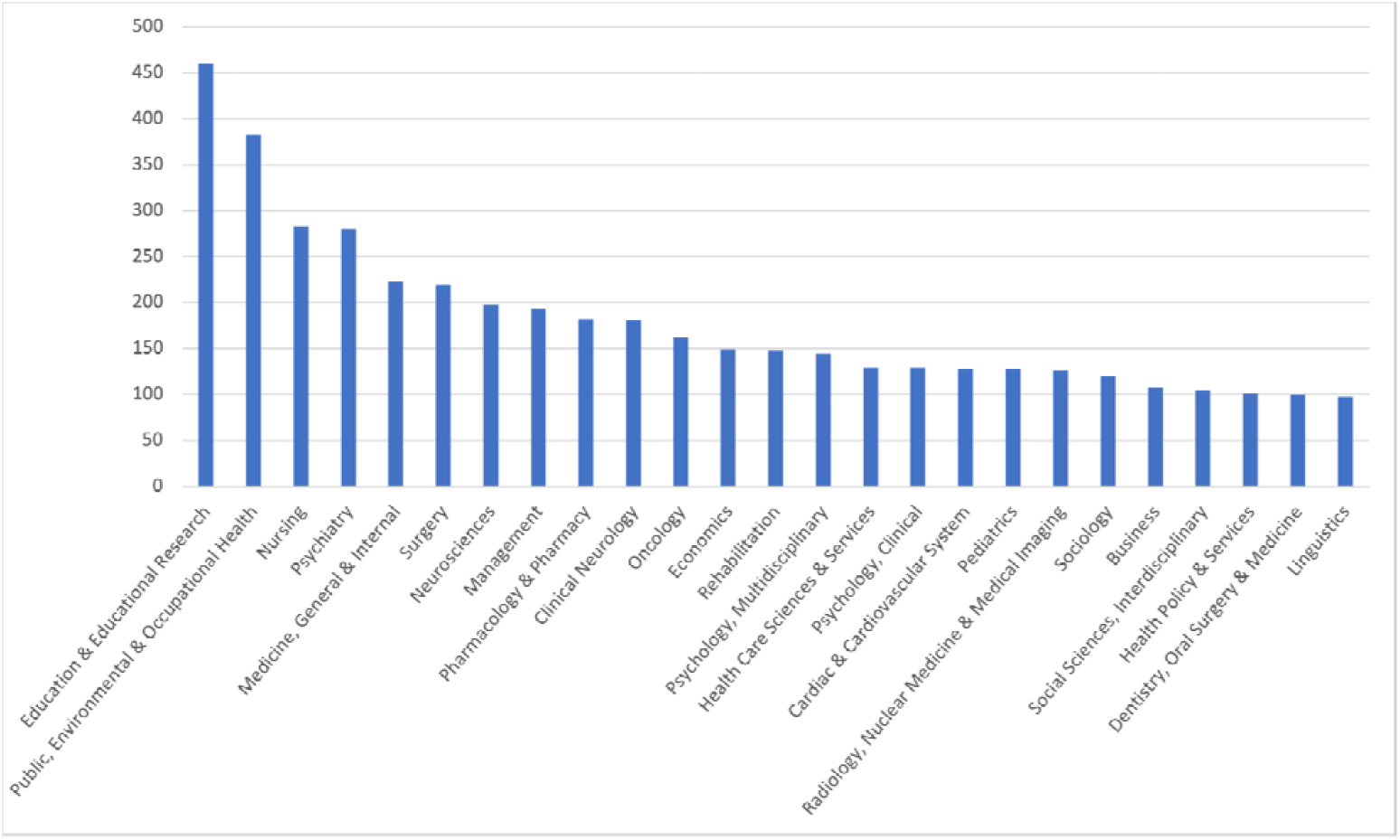
Top 25 Web of Science Categories based on the appearance of a journal

As most journals were cited multiple times (e.g., *Medical Education* was cited 50,372 times), we also calculated the count of categories based on the number of times a journal was cited overall. This accounted for categories appearing a total of 845,737 times. In this case, the most prevalent categories were: *Education, Scientific Disciplines* (n=202,663; 24.0%); *Health Care Sciences & Services* (n=202,125; 23.9%); and *Medicine, General & Internal* (n=76,555; 9.1%). See Table 3 for the top 10 categories based on count of all references, which accounted for 77% of all citations. Clinically-focused journals accounted for 159,334 (32.3%) of cited references. Figure 2 provides a depiction of the top 25 categories based on total number of appearances.

**Table 3:** Top 10 Web of Science (WoS) categories based on count of references

| Categories | Count of references (n; %) | Web of Science index and abridged scope note coverage |
| --- | --- | --- |
| Education, Scientific Disciplines | 202,663; 24.0 | SCIE: Education resources in the scientific disciplines, including biology, pharmacy, biochemistry, engineering, chemistry, nutrition, and medicine. |
| Health Care Sciences & Services | 202,125; 23.9 | SCIE: Resources on health services, hospital administration, health care management, health care financing, health policy and planning, health economics, health education, history of medicine, and palliative care. |
| Medicine, General & Internal | 76,555; 9.1 | SCIE: Resources on medical specialties such as general medicine, internal medicine, clinical physiology, pain management, military and hospital medicine. Resources focusing on family medicine and primary health care services are placed in the Primary Health Care category. |
| Education & Educational | 41,524; 4.9 | SSCI: Resources on the full spectrum of education, from theoretical to applied, from nursery school to Ph.D. Included in this category are resources on pedagogy and methodology as well as on |
| Research |  | the history of education, reading, curriculum studies, education policy, and the sociology and economics of education, as well as the use of computers in the classroom. |
| Surgery | 30,029; 3.6 | SCIE: Resources on general surgical topics including the different types of surgery (cardiovascular, neurosurgery, orthopedic, pediatric, or vascular); allied disciplines of surgery (surgical oncology, pathology, or radiology); and surgical techniques (arthroscopy, microscopy, or endoscopy). |
| Public, Environmental & Occupational Health | 28,596; 3.4 | SCIE: Resources dealing with epidemiology, hygiene, and health; parasitic diseases and parasitology; tropical medicine; industrial medicine; occupational medicine; infection control; and preventive medicine. Also included are resources on environmental health; cancer causes and control; aviation, aerosol, and wilderness medicine.<br>SSCI: Resources on social medicine, health behavior, health education, safety research, and community mental health. Resources concerned with the health of particular groups such as adolescents, elderly, or women are included in this category. |
| Nursing | 25,933; 3.1 | SCIE/SSCI: Resources on all aspects of nursing science and practice such as administration, economics, management, education, technological applications and all clinical care specialties. |
| Health Policy & Services | 14,536; 1.7 | SSCI: Resources on healthcare systems, including healthcare provision and management, financial analysis, healthcare ethics, health policy, and quality of care. |
| Psychiatry | 10,972; 1.3 | SCIE: Resources on clinical, therapeutic, research, and community aspects of human mental, emotional, and behavioral disorders.<br>SSCI: Resources that focus on the origins, diagnosis, and treatment of mental, emotional, or behavioral disorders. Areas covered in this category include adolescent and child psychiatry, forensic psychiatry, geriatric psychiatry, hypnosis, psychiatric nursing, psychiatric rehabilitation, psychosomatic research, and stress medicine. |
| Primary Health Care | 9,722; 1.1 | SCIE: Resources on all aspects of family medicine and primary health care services, including first contact, health assessments, laboratory and diagnostic procedures, medication management, disease prevention, early diagnosis and treatment and comprehensive strategies to improve the health status of individuals and communities. |
| Psychology, Multidisciplinary | 9,201; 1.1 | SSCI: Resources with a general or interdisciplinary approach to the field. Resources on philosophical psychology, psychobiology, and the history of psychology are included in this category. |
| Total | 651,856; 77.1 |  |

**Figure 2:**
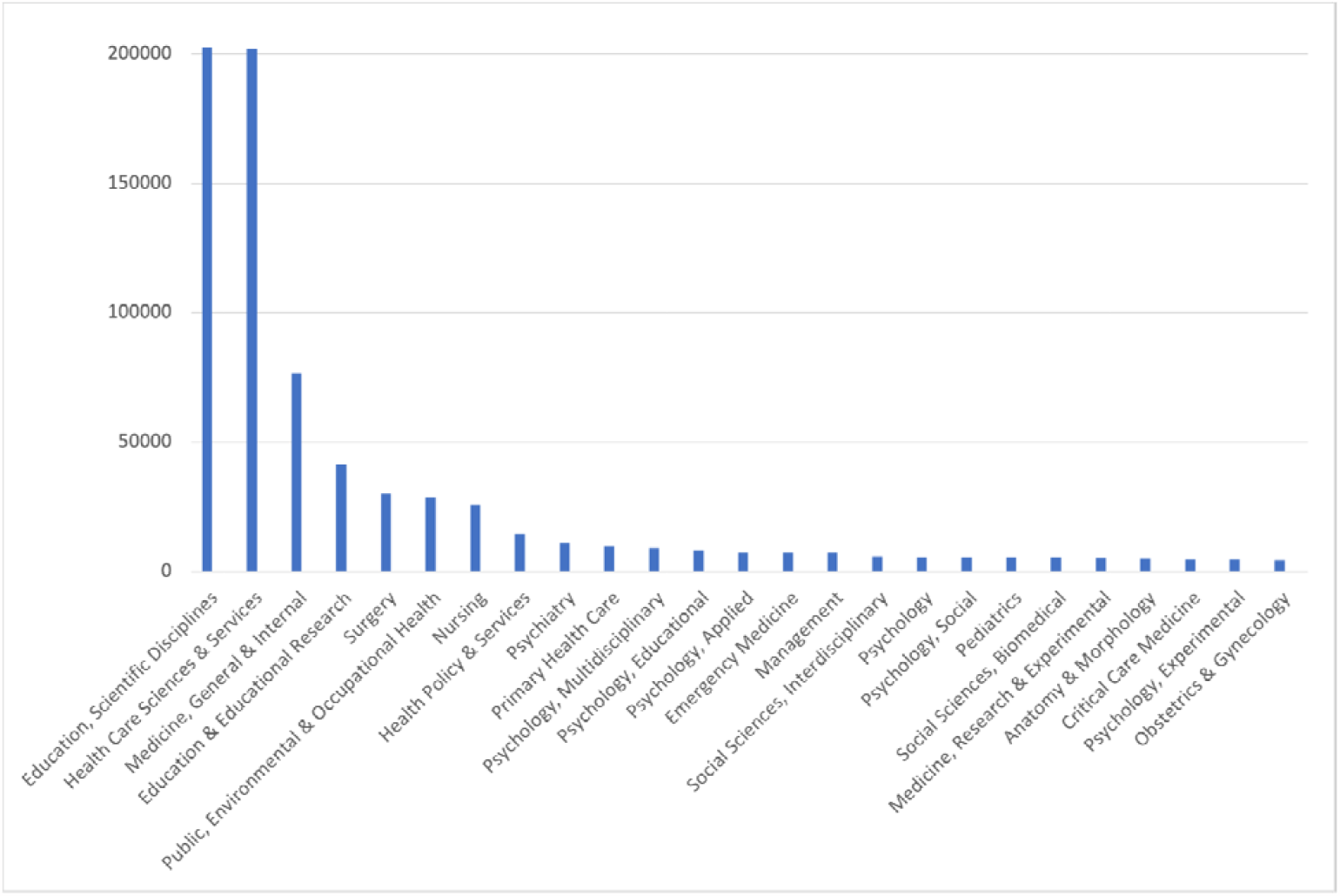
Top 25 Web of Science Categories by the total number of cited reference appearances

During the study time period (2001-2020), we observed consistent growth in the number of articles published in the MEJ-24, journals cited, references cited, and categories present. This growth is especially prevalent in the last five years. For example, while the overall number of citations to journals was 493,969, 49.2% of those citations occurred during the last 5 years of the time period analyzed (i.e., 2016-2020; see Table 4). The number of categories represented also increased over time, but at a slower rate. For example, from 2001-2005, there were 182 categories represented; where from 2016-2020, there were 233 categories represented, which includes over 90% of all WoS categories.

**Table 4:** Counts of journals, articles published, cited references, Web of Science categories represented and the sum of appearances in medical education journals between 2001-2020

| Year Span | Articles Published in MEJ-24 journals; % | Journals cited; % | Cited References; % | Web of Science Categories Present; % * |
| --- | --- | --- | --- | --- |
| 2001 - 2005 | 3885; 10.7 | 1365; 20.6 | 57155; 6.8 | 182; 71.1 |
| 2006 - 2010 | 6447; 17.8 | 2584; 39.0 | 121225; 14.3 | 207; 80.1 |
| 2011 - 2015 | 10631; 29.3 | 3955; 59.8 | 251452; 29.7 | 215; 83.9 |
| 2016 - 2020 | 15330; 42.2 | 5809; 87.8 | 415894; 49.2 | 233; 91.0 |
| Total | 36293; 100 | 6618; 100 | 493,969; 100 | 242; 94.5 |

\*Denotes the percentage of Web of Science categories (n=256)

When considering categories over time, the categories education, scientific disciplines, and healthcare sciences and services have continued to represent close to half of all categories represented. However, categories such as education and educational research and nursing have consistently demonstrated growth in representation. For example, between 2001-2005 education and educational research represented 2.28%, whereas during 2016-2020 this category represented 5.82% (See Figure 3).

**Figure 3:**
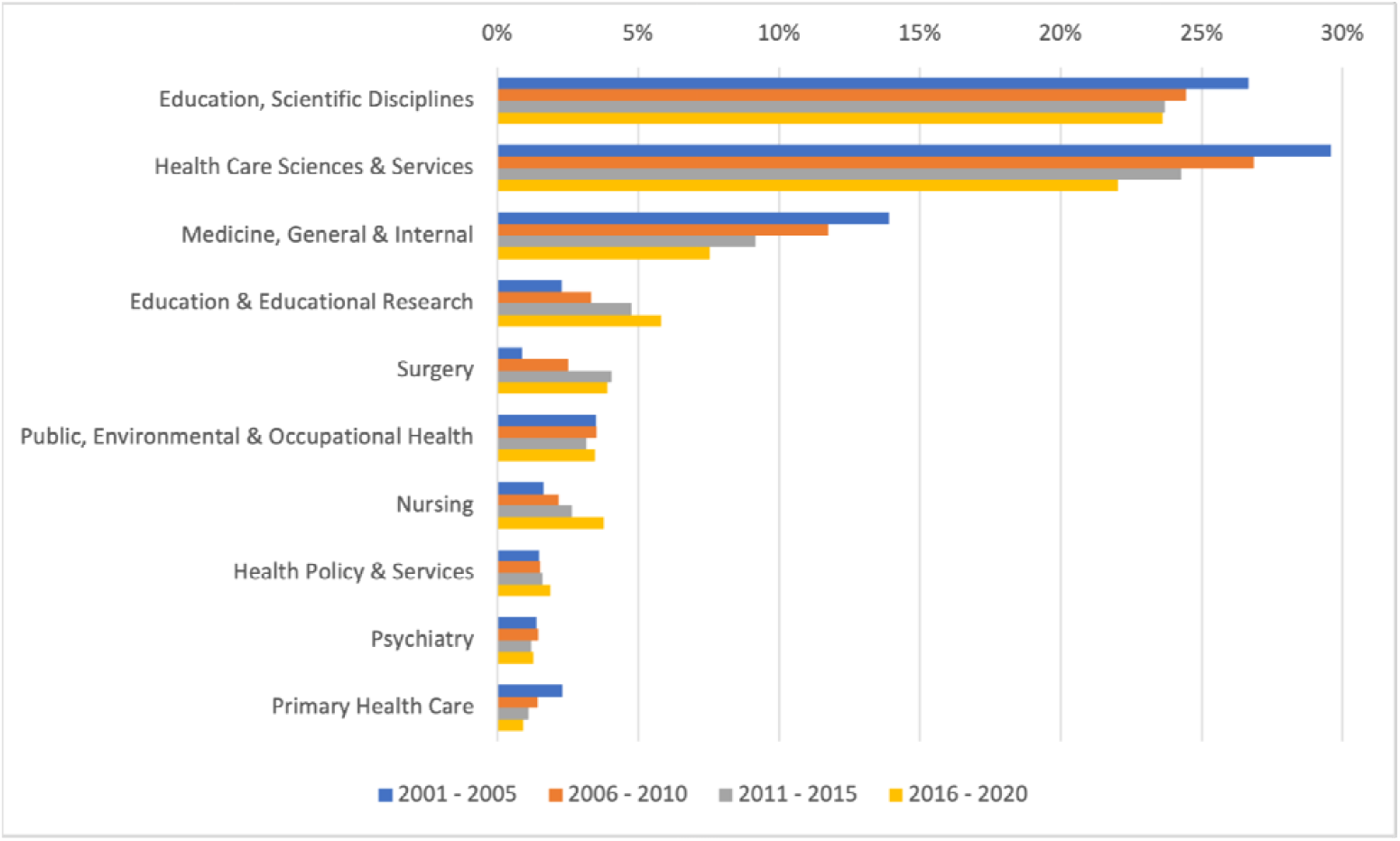
Top 10 Web of Science categories over time by percentage

## Discussion

Interdisciplinary research has been hailed as critical for helping to solve wicked problems [4], including those found in medical education. However, in recent studies Albert and colleagues questioned the interdisciplinarity of the field. They concluded that there is no convincing empirical evidence of interdisciplinarity, noting that the field’s research is inwardly focused [8, 9]. Based on our findings, which take into account a broader sample and categorization of the literature, we propose that the answer is not that simple. In some respects, medical education research could be labeled as inwardly focused. Yet, the growth in categories over time and the ongoing increased diversity of included categories suggests that this is not the whole story. In the paragraphs that follow, we consider our findings in relation to Albert and colleagues’ work and highlight some of the nuances now visible through our expanded approach. We also propose to the medical education research community some considerations if interdisciplinarity is to be a priority for the field.

As a conceptual replication, we believe that our study, to a degree, affirms Albert and colleague’s conclusions, but that it also extends and further deepens the level of evidence that can be brought to bear on the question of interdisciplinarity. In the earlier study, which utilized six author-defined clusters as compared to the 242 categories in this present study, Albert found that medical education research primarily cited references in the Applied Health and Medical Education clusters. These two clusters accounted for slightly over 80% of cited references. While in our study we do not have an exact match for these two clusters, we propose that the categories of *education, scientific disciplines* and *healthcare science and services* are quite similar to Albert et al.’s categories. These two categories represent nearly half (47.9%) of the citing references, such that our findings demonstrate that medical education research rests somewhat heavily on these two pillars of knowledge. Additionally, 40% of cited references appeared in journals in the MEJ-24 [19], which are designated specifically as medical education. This finding suggests that medical education research heavily cites its own research. Lastly, an inward focus is further demonstrated by our finding that most cited journals were contained in the science citation expanded index versus social sciences and arts and humanities indexes.

Our study extends Alberts and colleagues’ work by utilizing a much expanded set of categories and by examining medical education references over two decades versus a single time point. We propose that this expansion sheds light on nuances that suggest interdisciplinarity. For example, we observed that the number of categories represented increased over the time period studies, rising from 182 categories in 2001-2005 to 242 categories in 2016-2020 (which encompasses almost all of the WoS categories). This finding indicates a level of interdisciplinarity that is increasing and aligns with Laraviere’s research that over time authors across multiple fields have increasingly cited articles from outside their discipline, thus becoming more interdisciplinary [12]. In addition, this study’s expanded use of categories provides insights into potential interdisciplinary trends in medical education research that were previously invisible. Figure 1 illustrates the presence of cited references hailing from journals categorized as Business, Economics, Linguistics, Management, and Sociology. For example, the citing of business, economics and management journals may provide evidence of the increasing complexity of healthcare systems and medical education researchers’ efforts to inform education with topics like leadership [23, 24] and healthcare cost consciousness [25, 26]. Additionally, the inclusion of linguistics may be a nod towards the rise in electronic medical records and the novel language analysis methods used to mine them [27, 28].

As Albert points out, medical education journals, centers, and graduate programs tout themselves as being interdisciplinary [8]. However, when taken together this current study and Albert et al’s work suggests that there is potential for the field to grow in this regard. We propose that if the field desires increased interdisciplinarity, we must consider our current practices and determine if they foster or hinder this goal. For example, several researchers have explored the reality of social scientists and humanities scholars working in faculties of medicine, including those focused on education research [29, 30]. These studies found that the scholars adapted their traditional research approaches and perceptions of academic success to align with the epistemic norms of their newly adopted field of medicine. While these findings imply an inhospitable climate for interdisciplinarity, they also raise awareness of an issue, which now visible, can be addressed (if desired). To that end, beyond the current research climate in medical education, it is also important to seek and highlight opportunities for fostering future interdisciplinarity. Within medical education, there has been considerable growth in the number of graduate programs [31,32]. These programs may provide opportunities to welcome those from a variety of fields into medical education, but also to expose learners with roots in medicine to the benefits of interdisciplinarity and encourage such approaches. Journal editors in the field also provide opportunities for interdisciplinarity by featuring publication types such as the Cross Cutting Edge [1] published by *Medical Education* and Eye Openers featured in *Perspectives on Medical Education* [34], which challenge authors to introduce the field to theory, research methods and methodologies, and ideas from other fields thereby increasing interdisciplinarity.

Our work has several important limitations. First, the study sample was constructed from the MEJ-24, which is a proposed seed set of journals that does not comprehensively capture all articles that may contain medical education content. Second, we focused only on references to journal articles; it is possible that if we were to characterize the other publication types (e.g., books, reports, websites) we may have found increased interdisciplinarity. However, these other publication types were a minority of cited references. Thirdly, we relied on indexing at the journal level, but that does not necessarily imply that each paper published within that journal falls inside those particular categories. Lastly, we characterized the field as interdisciplinary or not based on its publications. This is a single measure of interdisciplinarity. Future work could further study interdisciplinarity by examining things such as the characteristics of author teams, the makeup of faculty in HPE programs, and the composition of students who graduate from HPE programs, to name just a few of the potentially relevant factors to consider when examining a field’ interdisciplinarity.

## Ethical Approval

Reported as not applicable

## Data

None reported

## Disclaimer

The views expressed in this article are those of the authors and do not necessarily reflect the official policy or position of the Uniformed Services University of the Health Sciences, Henry M. Jackson Foundation, the Department of Defense, or the U.S. Government.

## References

1. O’Sullivan PS, Stoddard HA, Kalishman S. Collaborative research in medical education: a discussion of theory and practice. Med Educ. 2010;44(12):1175–84.

2. Albert M, Friesen F, Rowland P, Laberge S. Problematizing assumptions about interdisciplinary research: implications for health professions education research. Adv Health Sci Educ Theory Pract. 2020;25(3):755–67.

3. Van Noorden R. Interdisciplinary research by the numbers. Nature. 2015;525(7569):306–7.

4. Pohl C, Truffer B, Hirsch-Hadorn G. Addressing Wicked Problems through Transdisciplinary Research, in Frodeman R (ed.), The Oxford Handbook of Interdisciplinarity, 2nd edn. Oxford, UK: Oxford University Press; 2017.

5. Hawick L, Cleland J, Kitto S. Getting off the carousel: Exploring the wicked problem of curriculum reform. Perspect Med Educ. 2017;6(5):337–43.

6. Onyura B, Crann S, Tannenbaum D, Whittaker MK, Murdoch S, Freeman R. Is postgraduate leadership education a match for the wicked problems of health systems leadership? A critical systematic review. Perspect Med Educ. 2019;8(3):133–42.

7. Varpio L, Aschenbrener C, Bates J. Tackling wicked problems: how theories of agency can provide new insights. Med Educ. 2017 Apr;51(4):353–65.

8. Albert M, Rowland P, Friesen F, Laberge S. Interdisciplinarity in medical education research: myth and reality. Adv Health Sci Educ Theory Pract. 2020;25(5):1243–53.

9. Albert M, Rowland P, Friesen F, Laberge S. Barriers to cross-disciplinary knowledge flow: The case of medical education research. Perspect Med Educ. 2022;11(3):149–55.

10. National Academies of Sciences, Engineering, and Medicine. Facilitating Interdisciplinary Research. Washington, DC: The National Academies Press; 2005.

11. Ninkov A, Frank JR, Maggio LA. Bibliometrics: Methods for studying academic publishing. Perspect Med Educ. 2022;11(3):173–76.

12. Larivière V, Gingras Y. On the relationship between interdisciplinarity and scientific impact. Journal of the American Society for Information Science and Technology. 2010 Jan;61(1):126–31.

13. Maggio LA, Costello JA, Ninkov AB, Frank JR, Artino AR Jr. The voices of medical education scholarship: Describing the published landscape. Med Educ. 2023;57(3):280–89.

14. Katritsis DG. Journal Impact Factor: Widely Used, Misused and Abused. Arrhythm Electrophysiol Rev. 2019;8(3):153–55.

15. Simons K. The misused impact factor. Science. 2008;322(5899):165.

16. Picho K, Maggio LA, Artino AR Jr. Science: the slow march of accumulating evidence. Perspect Med Educ. 2016;5(6):350–53.

17. Nosek BA, Errington TM. Making sense of replications. Elife. 2017;6:e23383.

18. Derksen M, Morawski J. Kinds of Replication: Examining the Meanings of “Conceptual Replication” and “Direct Replication”. Perspect Psychol Sci. 2022;17(5):1490–505.

19. Maggio LA, Ninkov A, Frank JR, Costello JA, Artino AR Jr. Delineating the field of medical education: bibliometric research approach (es). Med Educ. 2022;56(4):387–94.

20. Costello JA, Maggio LA, Ninkov A, Frank J, Artino A Jr. “Looking in or out?: A bibliometric study of the interdisciplinarity of medical education research” -Supplemental Files (Version 1); 2023. [Data set]. Zenodo. Published March 21, 2023. Available at: doi:10.5281/zenodo.7692524

21. Li K, Rollins J, Yan E. Web of science use in published research and review papers 1997–2017: a selective, dynamic, cross-domain, content-based analysis. Scientometrics. 2018;115(1):1–20.

22. American Board of Medical Specialities Member Boards. Retrieved 2 Aug 2022. https://www.abms.org/member-boards/

23. Matsas B, Goralnick E, Bass M, Barnett E, Nagle B, Sullivan EE. Leadership development in U.S. undergraduate medical education: A scoping review of curricular content and competency frameworks. Acad Med. 2022;97:899–908.

24. Sultan N, Torti J, Haddara W, Inayat A, Inayat H, Lingard L. Leadership development in postgraduate medical education: a systematic review of the literature. Academic Medicine. 2019 Mar 1;94(3):440–9.

25. Ginzburg SB, Schwartz J, Deutsch S, Elkowitz DE, Lucito R, Hirsch JE. Using a problem/case-based learning program to increase first and second year medical students’ discussions of health care cost topics. Journal of Medical Education and Curricular Development. 2019 Dec;6:2382120519891178.

26. Tartaglia KM, Kman N, Ledford C. Medical student perceptions of cost-conscious care in an internal medicine clerkship: a thematic analysis. Journal of general internal medicine. 2015 Oct;30:1491–6.

27. Pennebaker JW, Boyd RL, Jordan K, Blackburn, K. The development and psychometric properties of LIWC2015. Austin, TX: University of Texas at Austin;2015.

28. Ondov B, Attal K, Demner-Fushman D. A survey of automated methods for biomedical text simplification. J Am Med Inform Assoc. 2022;29(11):1976–88.

29. Paradis E, Reeves S. Key trends in interprofessional research: A macrosociological analysis from 1970 to 2010. J Interprof Care. 2013;27(2):113–22.

30. Albert M, Paradis E, Kuper A. Interdisciplinary fantasy: Social scientists and humanities scholars working in faculties of medicine. Investigating interdisciplinary collaboration: Theory and practice across disciplines. 2017:84–103.

31. Tekian A. Doctoral programs in health professions education. Medical teacher. 2014 Jan 1;36(1):73–81.

32. Artino Jr AR, Cervero RM, DeZee KJ, Holmboe E, Durning SJ. Graduate programs in health professions education: preparing academic leaders for future challenges. J Grad Med Educ. 2018;10(2):119–22.

33. Eva KW. The cross-cutting edge: striving for symbiosis between medical education research and related disciplines. Med Educ. 2008;42(10):950–1.

34. Perspectives on Medical Education. Submission Guidelines: Submitting an Article Online. Retrieved 14 Mar 2023. https://pmejournal.org/about/submissions

